# DNAJC12 stabilizes activated phenylalanine hydroxylase and reduces the concentration of L-Phe needed for activation

**DOI:** 10.1101/2025.07.18.665471

**Authors:** Mary Dayne S. Tai, Trond-André Kråkenes, Gloria Gamiz-Arco, Christer F. Didriksen, Juha P. Kallio, Marte I. Flydal, Fernando Moro, Aurora Martinez

## Abstract

Phenylalanine hydroxylase (PAH) is a tetrahydrobiopterin (BH_4_)-dependent enzyme that converts L-phenylalanine (L-Phe) to L-tyrosine. PAH dysfunction leads to the accumulation of L-Phe in the blood (hyperphenylalaninemia; HPA), which may reach neurotoxic levels, resulting in phenylketonuria (PKU). PKU is associated with pathogenic variants in *PAH*, most causing misfolding and instability, leading to decreased levels of PAH protein and activity. Recently, variants in the class C J-domain protein DNAJC12 have also been associated with HPA in patients, demonstrating the importance of protein homeostasis regulation for proper PAH function. DNAJC12 and PAH have previously been reported to interact, but the molecular and structural mechanisms behind complex formation have remained unclear. In this work, we show that DNAJC12 binds to PAH, but presents higher affinity for its L-Phe activated form, which resembles the conformation of unliganded tyrosine hydroxylase, a structurally and functionally-related enzyme that also binds to DNAJC12. At saturation, four monomers of DNAJC12 bind and stabilize the PAH tetramer, protecting it from aggregation and lowering the L-Phe concentration necessary for substrate-induced activation, without affecting the interaction of the enzyme with its cofactor BH_4_. Importantly, DNAJC12 also stabilizes and delays the aggregation of the PKU-associated variant PAH-p.R261Q. Furthermore, L-Phe activated wild-type or variant PAH is required to stimulate Hsc70 ATPase activity.

**SIGNIFICANCE STATEMENT:** Deficiencies in the cochaperone DNAJC12 have recently been linked to hyperphenylalaninemia, dystonia and intellectual disabilities as DNAJC12 regulates the proteostasis of the aromatic amino acid hydroxylases, including phenylalanine hydroxylase (PAH). This study explores the mechanisms of the PAH:DNAJC12 interaction and examines the functional effects of their complex formation on PAH activity and stability. These findings enhance our understanding on the pathogenic mechanisms behind *DNAJC12* variants and provide insights that could guide the development of drugs targeting this protein-protein interaction.

## INTRODUCTION

Hyperphenylalaninemia (HPA), with phenylketonuria (PKU; MIM261600) being its most severe manifestation, is an autosomal recessive inborn error of metabolism predominantly caused by genetic variants in the *PAH* gene (NM_000277.2), which encodes for the enzyme phenylalanine hydroxylase (PAH; EC 1.14.16.1). PAH converts L-phenylalanine (L-Phe) to L-tyrosine (L-Tyr) through a reaction that is dependent on non-heme iron (Fe^2+^), molecular oxygen (O_2_) and the enzymatic cofactor (6R)-L-erythro-5,6,7,8-tetrahydrobiopterin (BH_4_)^1,2^. Variants in *PAH* lead to deficient enzyme activity or reduced protein levels, subsequently causing an accumulation of L-Phe in the blood and brain. Late diagnosis of patients with HPA can lead to a broad phenotypic spectrum that includes mild autistic traits, hyperactive behavior, intellectual disabilities, dystonia and parkinsonism^3^.

PAH is a homotetrameric hepatic enzyme, where each 51-kDa subunit consists of an N-terminal regulatory domain (RD; residues 1-110), a catalytic domain (CD; residues 111-410) that facilitates the hydroxylation of L-Phe, and a C-terminal oligomerization domain (OD; residues 411-452)^4^. In response to increasing concentrations of L-Phe, full-length tetrameric PAH shows positive cooperativity^5,6^. The conformational transition induced by L-Phe is slow^7^ and leads to the dimerization of the RDs on opposite sides of the CD and ODs^8,9^, with L-Phe binding at the dimerization interface^10^.

Currently, more than 3000 PAH variants are registered in the BIOPKU database (http://www.biopku.org/). Most PKU-causing variants are missense (33.7%), deletions and splicing alterations and, recently, most missense variants have been classified using structural analysis and phenotypic data^11^. The major pathogenic mechanism associated with PKU variants is their instability, leading to decreased activity and increased degradation^12-14^, and PKU is traditionally viewed as a loss-of-function disease. Nevertheless, the characterization of a mouse model harboring one of the most common PKU-associated variants, PAH-p.R261Q, reveals a gain-of-function pathogenic mechanism, where the unstable and misfolded PAH protein forms toxic amyloid-like aggregates^15^.

Although most patients with HPA have variants in *PAH*, approximately 2% of cases are related to deficiencies in the synthesis or regeneration of the cofactor BH_416_. An even smaller proportion of HPA cases have more recently been attributed to biallelic mutations in the gene encoding the cochaperone DNAJC12 in patients without any variants in *PAH* or other genes related to BH_4_ synthesis or regeneration^17-19^. DNAJC12 is a specific interaction partner of the BH_4_-dependent aromatic amino acid hydroxylases (AAAH), including PAH, tyrosine hydroxylase (TH), and tryptophan hydroxylases 1 and 2 (TPH1 and TPH2)^20-22^, and thus DNAJC12 deficiency leads to reduced levels of functional AAAHs^17,23^. DNAJC12 is a class C J-domain containing protein (JDP), that recognizes and binds to specific clients and presents them to Hsp70 for the regulation of client protein homeostasis^24^. Furthermore, an Hsp70-independent holdase activity has recently been suggested for DNAJC12 towards its client TH, where the cochaperone increased the stability of TH and prevented its aggregation over time in vitro^21^.

DNAJC12 is a monomeric protein of 198 amino acids comprising a J-domain (JD; residues 14-76) that is essential for its interaction with Hsp70, a long linker region that includes a linker-helix (residues 87-100), and a highly conserved C-terminal domain (CTD; residues 176-198), which as demonstrated for TH is involved in client binding^21^. DNAJC12 recognizes and binds to the dimerized RDs of TH^21^ and is expected to recognize the same domain in the other AAAHs, despite the RD being the least conserved domain within this family of enzymes. Here, we report that DNAJC12 binds to PAH, preferentially to its allosterically L-Phe activated state, characterized by RD dimerization. Despite sequence divergence in the RDs of AAAHs, DNAJC12 recognizes PAH by the CTD. This interaction stabilizes PAH, particularly the RD, and lowers the L-Phe concentration needed for enzyme activation without affecting the positive cooperativity for L-Phe or its interactions with BH_4_. DNAJC12 also binds the L-Phe activated state of the variant PAH-p.R261Q, increases its activity at lower L-Phe concentrations, and delays aggregate formation in vitro, highlighting the PAH:DNAJC12 interaction as a potential therapeutic target for HPA and PKU treatment.

## RESULTS

### Four DNAJC12 monomers bind to the native PAH tetramer in the presence of L-Phe

The TH:DNAJC12 complex that includes two DNAJC12 molecules per TH tetramer, was previously obtained by combining TH and DNAJC12 in a 4:4 ratio, followed by analysis and purification using size exclusion chromatography (SEC)^21^. Unlike TH, which fully forms the complex under these conditions, PAH does not, and only a small proportion of PAH shows a shifted elution profile, indicating the presence of some PAH:DNAJC12 complex (Fig. 1A; left).

**Figure 1.**
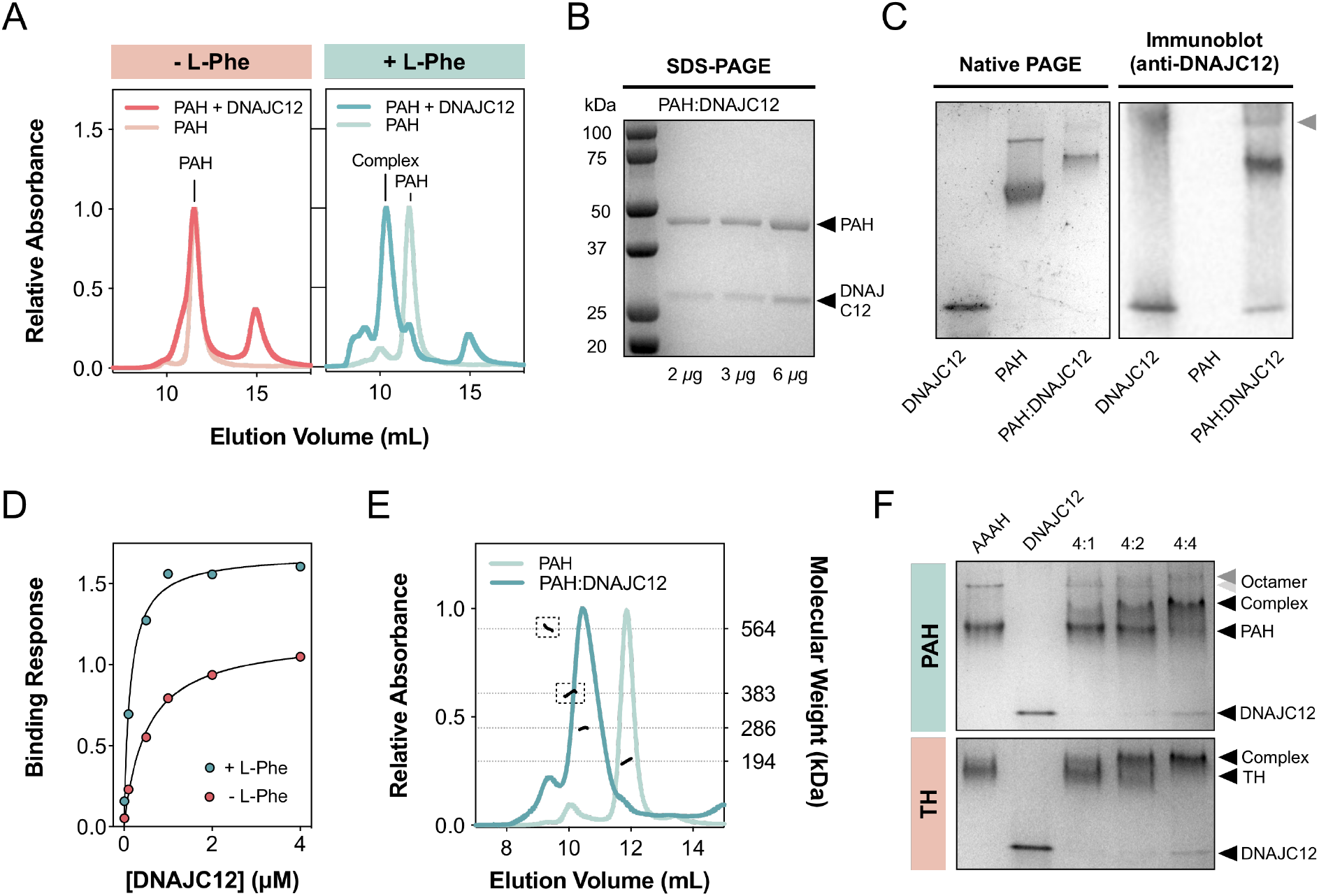
DNAJC12 binds to the activated form of PAH at 4(PAH subunit):4(DNAJC12) stoichiometry. **A. SEC chromatograms of PAH alone and with DNAJC12, with and without L-Phe**. The presence of DNAJC12 (20 µM) shifts the elution of a small proportion of tetrameric PAH (20 µM subunit), indicating complex formation, but PAH mostly remains unbound. In the presence of 1 mM L-Phe, DNAJC12 (20 µM) up-shifts the elution of tetrameric PAH (20 µM subunit), indicating efficient complex formation. **B. SDS-PAGE analysis of the PAH:DNAJC12 complex**. SDS-PAGE of the SEC purified complex (2, 3 or 6 µg protein), showing the co-elution of DNAJC12 (∼27 kDa) and PAH (∼48 kDa). **C. Native PAGE (left) and immunoblot (right) analyses of the PAH:DNAJC12 complex and controls**. The purified PAH:DNAJC12 complex migrates slower than PAH alone through the native gel (left). Using immunodetection with DNAJC12 antibodies (right) DNAJC12 is identified in the complex. A small proportion of the sample is detected as part of a larger complex, likely to be the octameric PAH:DNAJC12 complex (gray arrow; see below 1F). **D. BLI analyses of full-length DNAJC12-concentration dependent binding to PAH, with or without L-Phe**. L-Phe increases the affinity of DNAJC12 for PAH by more than 3-fold (K_D_=139 ± 19 nM with L-Phe; blue symbols; and K_D_=508 ± 59 nM without L-Phe; red symbols). **E. Determination of PAH:DNAJC12 stoichiometry by SEC-MALS**. Analysis of the PAH:DNAJC12 complex (200 µg; dark blue) and PAH (200 µg; light blue) by SEC-MALS provided an estimated size of 286.2 ± 1.8 kDa for the complex, consistent with the binding of four DNAJC12 monomers to each PAH tetramer (194.1 ± 4.6 kDa). In addition, octameric PAH (383.1 ± 5.4 kDa) is also detected and forms a complex with 8 DNAJC12 monomers (564.0 ± 4.6 kDa) (stippled boxes). **F. Native PAGE analyses of PAH and TH with varying amounts of DNAJC12**. Addition of varying amounts of DNAJC12 (4:1, 4:2 and 4:4 AAAH subunit:DNAJC12 ratio) to a constant amount of PAH or TH, show that a proportion of PAH remains unbound even at 4:4 ratio, while all TH has bound at the same conditions. A larger PAH oligomer (light gray arrow) is also detected and also increases in size, consistent with the formation of the octameric PAH:DNAJC12 complex (dark gray arrow).

While PAH and TH share sequence and structural conservation to a large degree, the unliganded states of both proteins display different structural features in the resting state. In particular, the PAH-RDs are monomeric^4,8^, while the TH RDs form dimers on the opposite sides of the CD+OD^25^. However, in the presence of L-Phe, the PAH-RDs dimerize^9,10^. As DNAJC12 was recently described to bind around the dimeric RDs of TH^21^, we hypothesized that the large conformational change associated to enzyme activation by L-Phe, leading to dimerization of the RDs, could improve the binding of DNAJC12 to PAH. Thus, we repeated the binding experiment supplementing the running and sample buffers with 1 mM L-Phe, a concentration that activates human PAH^5^. In the presence of L-Phe, a large up-shift in the elution profile of PAH was observed when DNAJC12 was added (Fig. 1A; right), indicating the efficient formation of the PAH:DNAJC12 complex. The presence of the complex in this fraction was demonstrated by PAGE in denaturing and non-denaturing conditions. SDS-PAGE analysis revealed two bands corresponding to DNAJC12 (23.5 kDa) and PAH (52 kDa), confirming the co-elution of the two proteins (Fig. 1B), and analysis of the same sample using native PAGE showed slower migration through the polyacrylamide matrix, as compared to PAH alone (Fig. 1C; left). Moreover, subsequent immunoblotting at native conditions using antibodies against DNAJC12 detected the protein in the DNAJC12 control sample and co-migration of PAH and DNAJC12 in the expected band for the PAH:DNAJC12 complex, but not in the PAH control, supporting stable complex formation (Fig. 1C; right).

The effect of L-Phe on the affinity between PAH and DNAJC12 was then investigated by bio-layer interferometry (BLI). The binding response between immobilized biotinylated PAH on streptavidin biosensors and varying DNAJC12 concentrations were measured with and without 1 mM L-Phe. In agreement with the SEC results (Fig. 1A), we found that L-Phe increases the affinity of DNAJC12 for PAH by more than 3-fold (K_D_=139 ± 19 nM) than without L-Phe (K_D_=508 ± 59 nM) (Fig. 1D). Furthermore, the stoichiometry of the PAH:DNAJC12 complex was determined by SEC coupled with multi-angle light scattering (SEC-MALS). While PAH alone presented the expected mass for the tetrameric enzyme (194.1 ± 4.6 kDa), the complex was estimated to be 286.2 ± 1.8 kDa, ∼92 kDa larger than PAH alone (Fig. 1E). As each DNAJC12 monomer is 23.5 kDa, these results indicate that four DNAJC12 molecules bind to each PAH tetramer, unlike TH that only binds two DNAJC12s per tetramer. In accordance with these results, we also analyzed samples containing PAH or TH with increasing amounts of DNAJC12 (4:1, 4:2 or 4:4) by native PAGE (Fig. 1F). In both cases, the complexes migrated slower through the gel than their unbound counterparts, but in the PAH samples combined with DNAJC12 at a 4:4 PAH:DNAJC12 ratio, unbound PAH still remained, differently to TH, for which full complex formation had already been achieved at the same conditions (Fig. 1F). While TH and activated PAH share structural similarities, it is evident that DNAJC12 binds to both proteins at different stoichiometries.

It is also worth noting that a small proportion of PAH exists as larger oligomers, which by SEC-MALS were determined to have a molecular size of 383.1 ± 5.4 kDa (Fig. 1E), in agreement with previous reports of stable PAH octamers, especially in the presence of L-Phe^26^. Interestingly, this octameric species also seems to bind DNAJC12, as an even larger species (564.0 ± 4.6 kDa; ∼180 kDa larger than the experimentally determined size of octameric PAH), corresponding to the binding of eight DNAJC12 monomers to each PAH octamer (Fig. 1E; stippled boxes). Evidence of this octameric species and the complex that it forms with DNAJC12 is also shown in the native gels and immunoblotting results as species that migrate through the polyacrylamide matrix even slower than the more abundant tetrameric PAH:DNAJC12 complex (Figs. 1C and 1F).

### DNAJC12 stabilizes PAH by binding to the PAH-RDs through its C-terminal client binding domain (CTD)

The DNAJC12 variant c.524G>A (p.W175Ter, also referred to as DNAJC12(1-174)) lacks the C-terminal 23 amino acids^23^ that includes the evolutionarily-conserved heptapeptide sequence ^192^KFRNYEI^198^. This is the most common disease-associated DNAJC12 variant identified to date^27^ and has recently been shown to be unable to bind to TH, enabling the identification of this region (also known as the C-terminal client-binding domain (CTD); residues 176-198) as an essential determinant for TH:DNAJC12 complex formation^21^. To investigate whether DNAJC12 also binds to PAH using the same region, we purified the truncated DNAJC12 variants DNAJC12(1-174) and DNAJC12(1-190), and tested whether they could bind to PAH using the same SEC binding assay where the buffer was supplemented with 1 mM L-Phe (Fig. 2A). In contrast to the results obtained with full-length DNAJC12 (Fig. 1A; blue figure), both truncated DNAJC12 variants do not alter the elution of PAH, indicating the necessity of the CTD for interaction with PAH (Fig. 2A). Furthermore, to probe whether DNAJC12 binds to the PAH-RD, as observed for TH^21^, we purified the PAH RD and CD+OD separately and analyzed binding by SEC. Due to the propensity of PAH-RD to aggregate^10^, we resorted to using the PAH(RD) or PAH(CD+OD) fused to a maltose binding protein (MBP) tag for the SEC binding experiments (Fig. 2B). The presence of MBP maintains the stability of the PAH(RDs) and does not inhibit dimerization, as shown in the up-shifted elution of PAH(RD) in the presence of L-Phe compared with the control sample without L-Phe. A further up-shift in the elution of PAH(RD) was observed in the presence of DNAJC12 and L-Phe, indicating complex formation (Fig. 2B; top panel). On the other hand, the presence of DNAJC12 or L-Phe did not affect the elution profile of PAH(CD+OD), confirming that DNAJC12 recognizes the PAH-RD and not the CD+OD (Fig. 2B; bottom panel).

**Figure 2.**
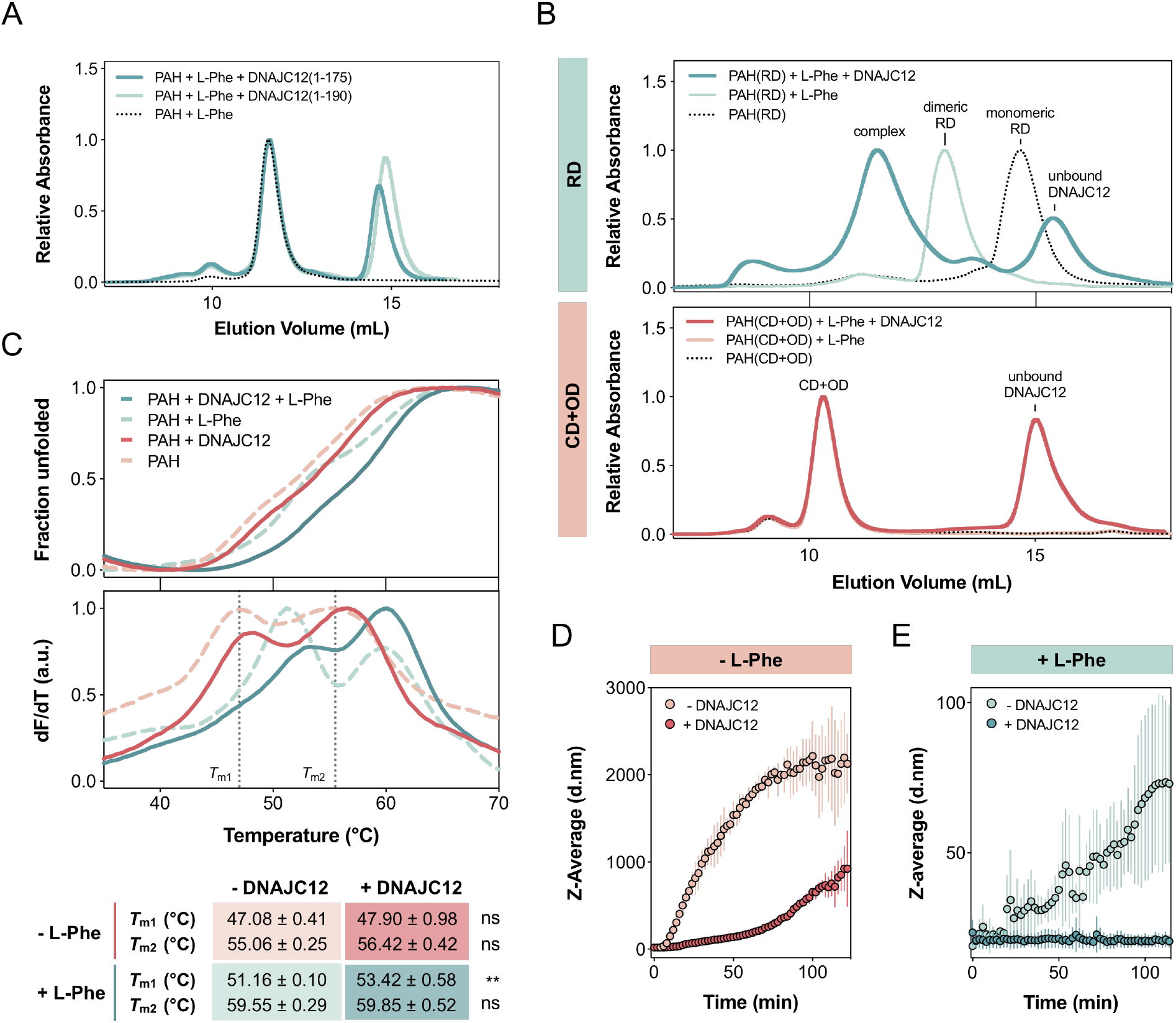
DNAJC12 binds and stabilizes the PAH-RD. **A. SEC chromatograms of PAH alone and with C-terminal truncated DNAJC12 variants, in the presence of 1 mM L-Phe**. DNAJC12(1-175) (dark blue; 40 µM) or DNAJC12(1-190) (light blue; 40 µM), do not affect the elution of PAH (stippled line; 20 µM subunit). **B. SEC chromatograms of MBP-PAH(RD) or MBP-PAH(CD+OD) alone and with 1 mM L-Phe; effect of DNAJC12 binding**. SEC analysis of MBP-PAH(RD) (20 µM subunit; top) or MBP-PAH(CD+OD) (20 µM subunit; bottom) in the presence (dark color line) or absence (light color line) of DNAJC12 (40 µM), without (stippled lines) or with 1 mM L-Phe (solid lines). **C. DSF-monitored unfolding**. Representative thermograms for PAH (1.92 µM subunit) alone (stippled and light-colored lines) or in presence of 3.84 μM DNAJC12 (solid and dark-colored lines) with (blue) or without 1 mM L-Phe (pink) show the biphasic transition of PAH unfolding, corresponding to the denaturation of the RD (*T*_m1_) and the CD+OD (*T*_m2_). L-Phe increases the thermal stability of both the RD and CD+OD, and DNAJC12 increases the thermal stability of the RD, without L-Phe. **D. and E. DLS-monitored aggregation of PAH in the presence and absence of DNAJC12, without** **(D) or with (E) 1 mM L-Phe**. Z-average hydrodynamic diameter (d.nm; mean ± SD; n=3 independent samples) of initially tetrameric PAH (10 μM subunit), for 120 min at 37 °C without (light symbols) and with DNAJC12 (10 μM; dark symbols) in the absence (red symbols) or presence of L-Phe (blue symbols).

We then tested whether DNAJC12 exerted a stabilizing effect on PAH upon their interaction. Using differential scanning fluorimetry (DSF), we monitored the thermal unfolding of PAH in the presence or absence of DNAJC12 with 1 mM L-Phe. PAH displays a biphasic transition (Fig. 2C) corresponding to the denaturation of the RD (*T*_m1_=47.08 ± 0.41 °C) and the CD+OD (*T*_m2_=55.06 ± 0.25 °C)^10,28^ and thus, it was possible to monitor the effect of DNAJC12 on these separate domains by comparing the obtained melting temperatures (*T*_m1_ and *T*_m2_) in the presence and absence of DNAJC12, with or without L-Phe. Corresponding well with previous reports that demonstrate the thermal stabilization of the PAH-RD induced by the presence of L-Phe, we observed a 4 °C increase in *T*_m1_, to 51.16 ± 0.10 °C with 1 mM L-Phe, which is related to its dimerization in response to substrate-induced activation^10^. A similar increase in *T*_m2_ was observed, indicating the stabilization of the CD+OD upon the addition of 1 mM L-Phe, consistent with the binding of L-Phe to the active site. In agreement with our previous SEC results showing that DNAJC12 binds to the PAH-RD and not the CD+OD (Fig. 2B), we found that the presence of DNAJC12 did not significantly increase the *T*_m2_ of PAH regardless of the presence or absence of L-Phe (Fig. 2C). Interestingly, however, a 2 °C increase in PAH *T*_m1_ was observed in the presence of DNAJC12 with 1 mM L-Phe (*T*_m1_=53.42 ± 0.58 °C) as compared to when the enzyme is alone (*T*_m1_=51.16 ± 0.10 °C), while no significant changes were observed when there was no L-Phe (*T*_m1_=47.08 ± 0.41 °C for PAH alone; *T*_m1_=47.90 ± 0.98 °C for PAH with DNAJC12), indicating that DNAJC12 binding results in the stabilization of the PAH-RDs, that occurs only in the presence of L-Phe.

To assess whether the thermal stabilization of PAH also resulted in a delay in its aggregation over time in vitro, we monitored changes in the particle size of PAH with and without DNAJC12, both in the absence and presence of 1 mM L-Phe, for 120 min. Consistent with the significant thermal stabilization imparted by L-Phe to PAH (Fig. 2C), we observed that the presence of the substrate significantly delays PAH aggregation in vitro (Figs. 2D and 2E). Additionally, DNAJC12 shows protection towards PAH aggregation without L-Phe (Fig. 2D), but is even more effective when L-Phe is present (Fig. 2E), which seems related to the increased affinity and saturation of DNAJC12 binding to the substrate activated PAH (Fig. 1A,D). These findings collectively highlight the stabilizing effect of DNAJC12 on PAH, particularly when combined with L-Phe.

### DNAJC12 decreases the L-Phe concentration required for enzyme activation

To investigate whether DNAJC12 binding affects the L-Phe dependent regulation of PAH and/or the enzyme kinetic parameters, we performed both PAH stability and activity assays with varying L-Phe concentrations. Because DNAJC12 binds to the PAH-RDs, which are stabilized upon L-Phe binding (Fig. 1C), we monitored the thermal unfolding of full-length PAH in the presence or absence of DNAJC12 using DSF at varying L-Phe concentrations and monitored any changes in *T*_m1_, corresponding to PAH-RD unfolding. Plotting the *T*_m1_ values recorded at varying L-Phe concentrations provided the substrate concentration at which half-maximal stabilization of this domain is reached (EC_50_). We found that the EC_50_ for PAH-RD alone (255.9 ± 24.4 µM) is approximately 10 times higher than in the presence of DNAJC12 (25.6 ± 4.3 µM) (Fig. 3A, top and bottom panels), indicating that DNAJC12 binding lowers the concentration of L-Phe required to stabilize the PAH-RDs.

**Figure 3.**
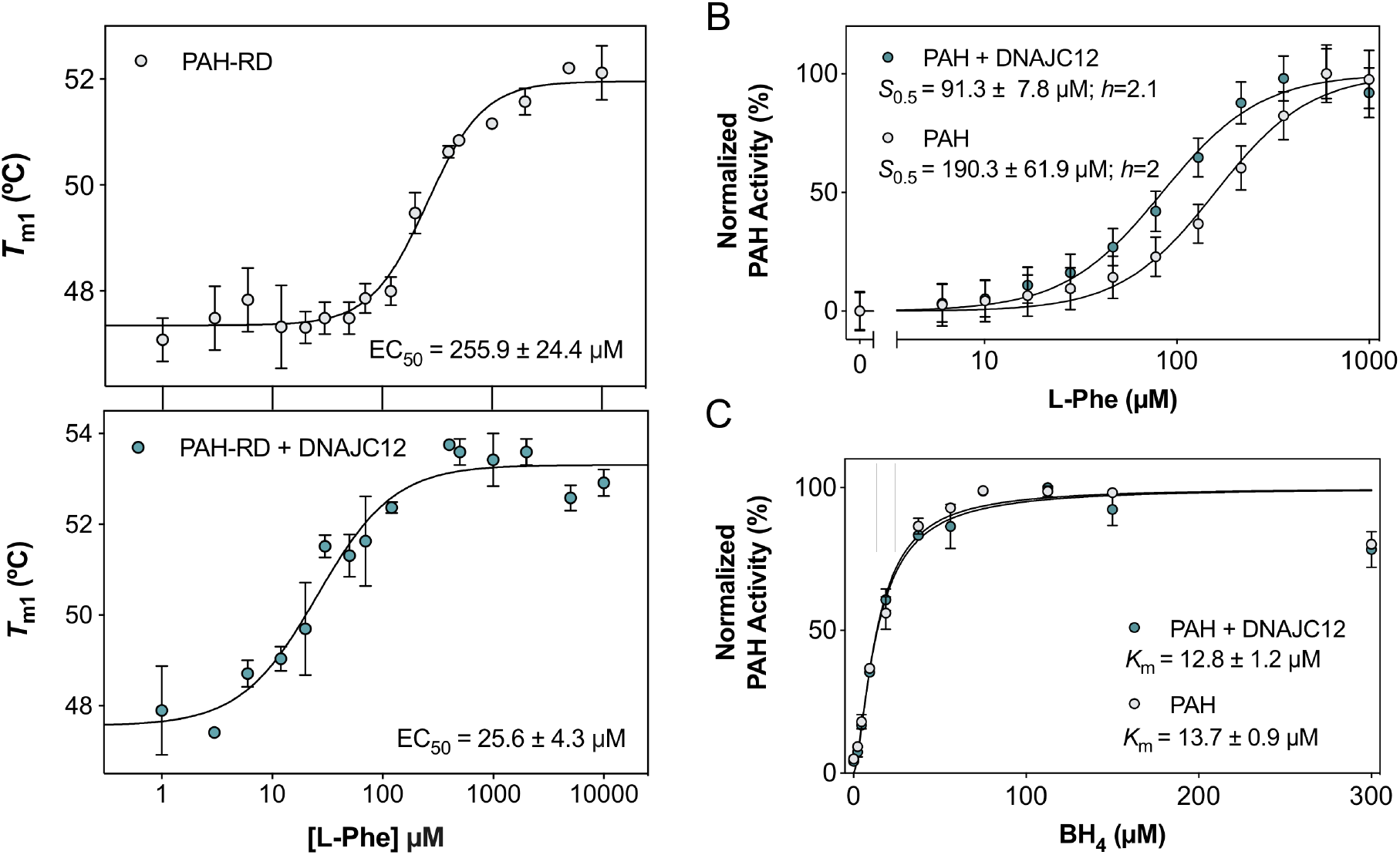
DNAJC12 lowers the L-Phe concentration necessary for substrate-induced activation. **A. Concentration-dependent stabilization of the PAH-RD by L-Phe without (upper panel) and with DNAJC12 (lower panel)**. The thermal unfolding of PAH-RD at varying L-Phe concentrations (0-10000 µM) with (bottom) and without (top) excess DNAJC12 (4:8 PAH:DNAJC12 subunit ratio) was monitored by following the first melting temperature transition (*T*_m1_) of PAH. Fitting the data to a four-parameter logistic nonlinear regression provides EC_50_ values (mean ± SD; n=3 independent samples). **B. and C. Enzymatic activity of PAH at variable L-Phe (0-1 mM; B) and BH**_**4**_ **concentrations (0-0.3 mM; B)**. The L-Phe concentration dependent activity of PAH presents positive cooperativity, with the indicated *S*_0.5_ and Hill coefficient (*h*). The presence of DNAJC12 maintains the positive cooperativity (*h*=2) but increases the affinity of PAH for L-Phe, as indicated by the 54% reduction in *S*_0.5_. **C)** The BH_4_ concentration dependence of PAH activity follows Michaelis-Menten kinetics, and DNAJC12 does not alter the *K*_m_ value for the cofactor. Activity data are presented as normalized results, n=3 independent experiments, each with technical triplicates, fitted to a four-parameter dose-response curve. *S*_0.5_ and *K*_m_ values are presented as mean ± SD.

To determine whether these results translate to a lower L-Phe concentration necessary to elicit the cooperative conformational changes during PAH activation, we measured PAH activity in the presence and absence of DNAJC12 at varying L-Phe concentrations. Consistent with previous studies, we found that PAH alone presents positive cooperativity for L-Phe providing half maximal activity (*S*_0.5_) at 190.3 ± 61.9 µM (Hill coefficient; *h*=2). Interestingly, in the presence of DNAJC12, a 54% decrease in PAH *S*_0.5_ to 91.3 ± 7.8 µM was obtained (Fig. 3B), without affecting the positive cooperativity (*h*=2.1), indicating that DNAJC12 increases PAH activity at lower substrate concentrations. In addition, PAH activity measurements at a range of BH_4_ concentrations show the hyperbolic dependence of the activity vs BH_4_ concentration and that DNAJC12 does not affect the *K*_m_-value for the cofactor (Fig. 3C). Together, these findings suggest that DNAJC12 lowers the L-Phe concentration needed to stabilize the dimerized PAH-RD conformation and to achieve maximal enzymatic activity, without affecting the affinity of PAH for BH_4_.

### DNAJC12 also binds and stabilizes the PKU-associated variant PAH-p.R261Q and delays its aggregation in vitro

We tested whether the misfolded and unstable PKU-associated variant PAH-p.R261Q can also be recognized by DNAJC12 as a client protein. Using SEC, we found that DNAJC12 is able to form a complex with PAH-p.R261Q in the presence of L-Phe (Fig. 4A). This variant has been found to form aggregates in a mouse model^15^, and thus, we tested if the stabilization imparted by DNAJC12 also translates to a delay in its aggregation. By monitoring any changes in PAH-p.R261Q particle size over time, without or with L-Phe or DNAJC12 using DLS, we observed that DNAJC12 delays the aggregation of PAH-p.R261Q, even in the absence of L-Phe, but with even better efficacy in the presence of L-Phe (Fig. 4B). In addition, we also observed a delay in PAH-p.R261Q aggregation in the presence of 1 mM L-Phe alone (Fig. 4B), showing that the variant responds in a similar manner to what is observed for wild-type (WT) PAH (Fig. 2D,E).

**Figure 4.**
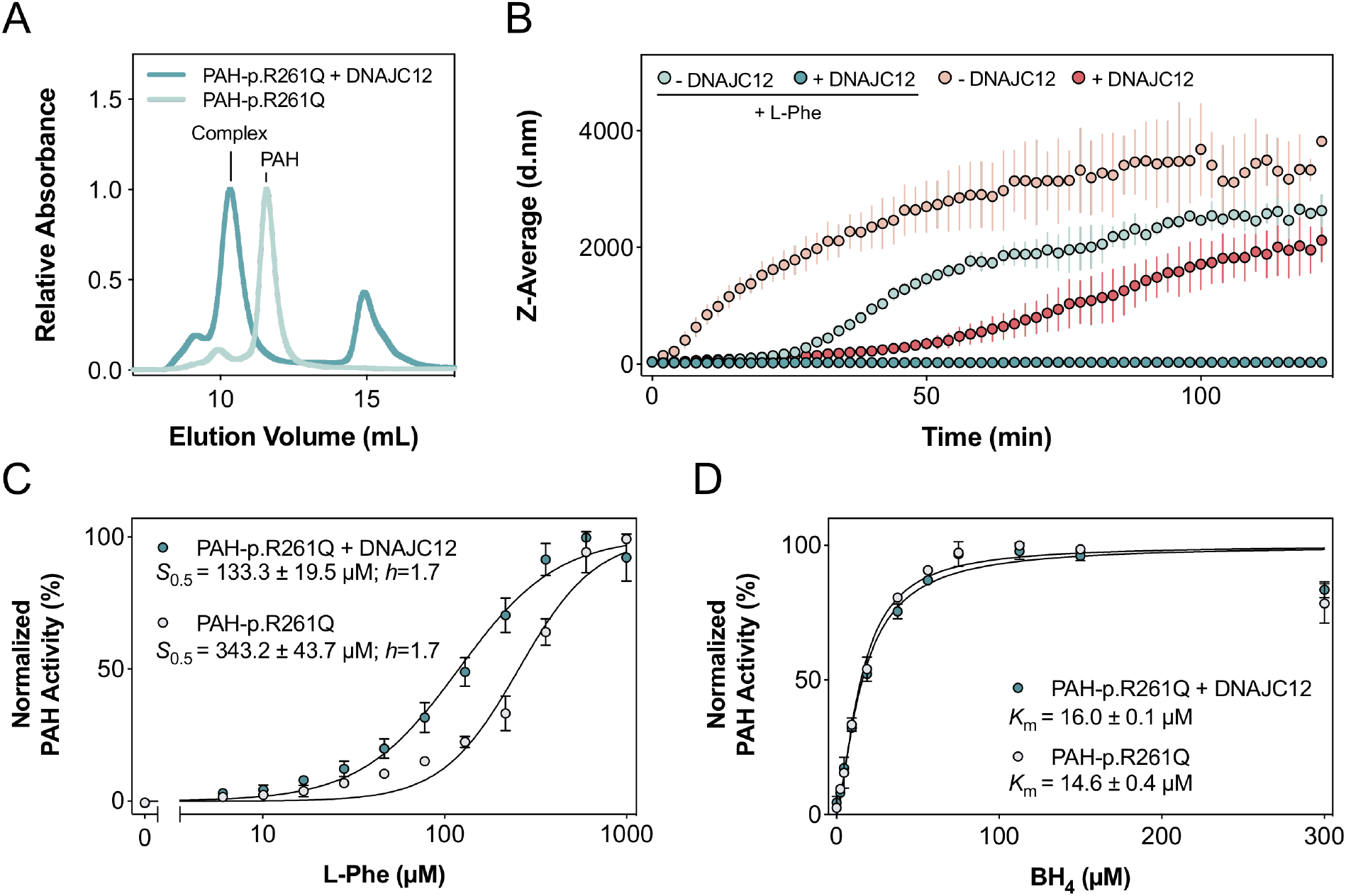
DNAJC12 binds and stabilizes the PKU-associated variant PAH-p.R261Q, and decreases the L-Phe concentration necessary for its enzymatic activation. **A. SEC chromatograms of PAH-p.R261Q alone and in the presence of DNAJC12**. Excess DNAJC12 (25 µM) shifts the elution of tetrameric PAH-p.R261Q (20 µM subunit) in the presence of 1 mM L-Phe, indicating efficient complex formation. **B. DLS-monitored aggregation of PAH-p.R261Q with and without DNAJC12 or L-Phe**. Z-average hydrodynamic diameter (d.nm; mean ± SD; n=3 independent samples) of initially tetrameric PAH-p.R261Q (10 μM subunit), for 120 min at 37 °C without (light symbols) and with DNAJC12 (10 μM; dark symbols), without (red symbols) or with 1 mM L-Phe (blue symbols). **C. and D. Enzymatic activity of PAH-p.R261Q at variable L-Phe (0-1 mM; C) and BH**_**4**_ **concentrations (0-0.3 mM; C)**. The L-Phe concentration dependent activity of PAH presents positive cooperativity, with the indicated *S*_0.5_ and Hill coefficient (*h*). The presence of DNAJC12 does not affect the *h*-value (*h*=1.7) but increases the affinity of PAH-p.R261Q for L-Phe, as indicated by the 65% reduction in *S*_0.5_. **D)**. The BH_4_ concentration dependence of PAH-p.R261Q activity follows Michaelis-Menten kinetics, and DNAJC12 does not alter the *K*_m_ value for the cofactor. Activity data are presented as normalized results, n=3 independent experiments, each with technical triplicates, fitted to a four-parameter dose-response curve. *S*_0.5_ and *K*_m_ values are presented as mean ± SD.

In order to determine whether DNAJC12 binding and consequent stabilization of PAH-p.R261Q also result in the decrease of L-Phe threshold for enzyme activation without any effect on BH_4_ regulation, we measured the conversion of L-Phe to L-Tyr by PAH-p.R261Q, in the presence and absence of DNAJC12, at varying concentrations of L-Phe or BH_4_. Consistent with previous studies that report that this variant also present positive cooperativity, with a somehow increased *S*_0.512_ compared with WT PAH, we measured a *S*_0.5_ = 343.2 ± 43.7 µM L-Phe (Fig. 4C), around double the L-Phe concentration needed for WT PAH, and a *h* = 1.7 (Fig. 3B). In the presence of DNAJC12, the *S*_0.5_ of PAH-p.R261Q decreases by 65% to 133.3 ± 19.5 µM (Fig. 4C). As with WT PAH, DNAJC12 also does not affect the *K*_m_-value of BH_4_ for PAH-p.R261Q (Fig. 4D).

### PAH and DNAJC12 synergistically activate Hsc70 ATPase activity with L-Phe

As a JDP, DNAJC12 is known to activate Hsc70 ATPase activity, though at much lower levels compared to canonical JDPs DNAJA2 and DNAJB1^21^(Fig. 5A), and as seen in Fig. 5B, L-Phe alone did not stimulate its activity. We investigated whether PAH and DNAJC12 could synergistically activate Hsc70 ATPase activity, with or without L-Phe. We found that PAH or PAH-p.R261Q alone do not stimulate Hsc70 ATPase activity, regardless of the presence of L-Phe (Fig. 5B), and the addition of PAH or PAH-p.R261Q only slightly improves DNAJC12 activity. However, the combination of either of these PAHs with L-Phe significantly enhances the stimulatory effect of DNAJC12 on Hsc70 ATPase activity (Fig. 5B), supporting the importance of L-Phe for PAH:DNAJC12 complex formation. This effect appears specific to DNAJC12, as the same was not observed with canonical JDPs DNAJA2 and DNAJB1, whose stimulatory effect on Hsc70 ATPase activity remains unaffected by PAH addition, even with L-Phe (Fig. 5A).

**Figure 5.**
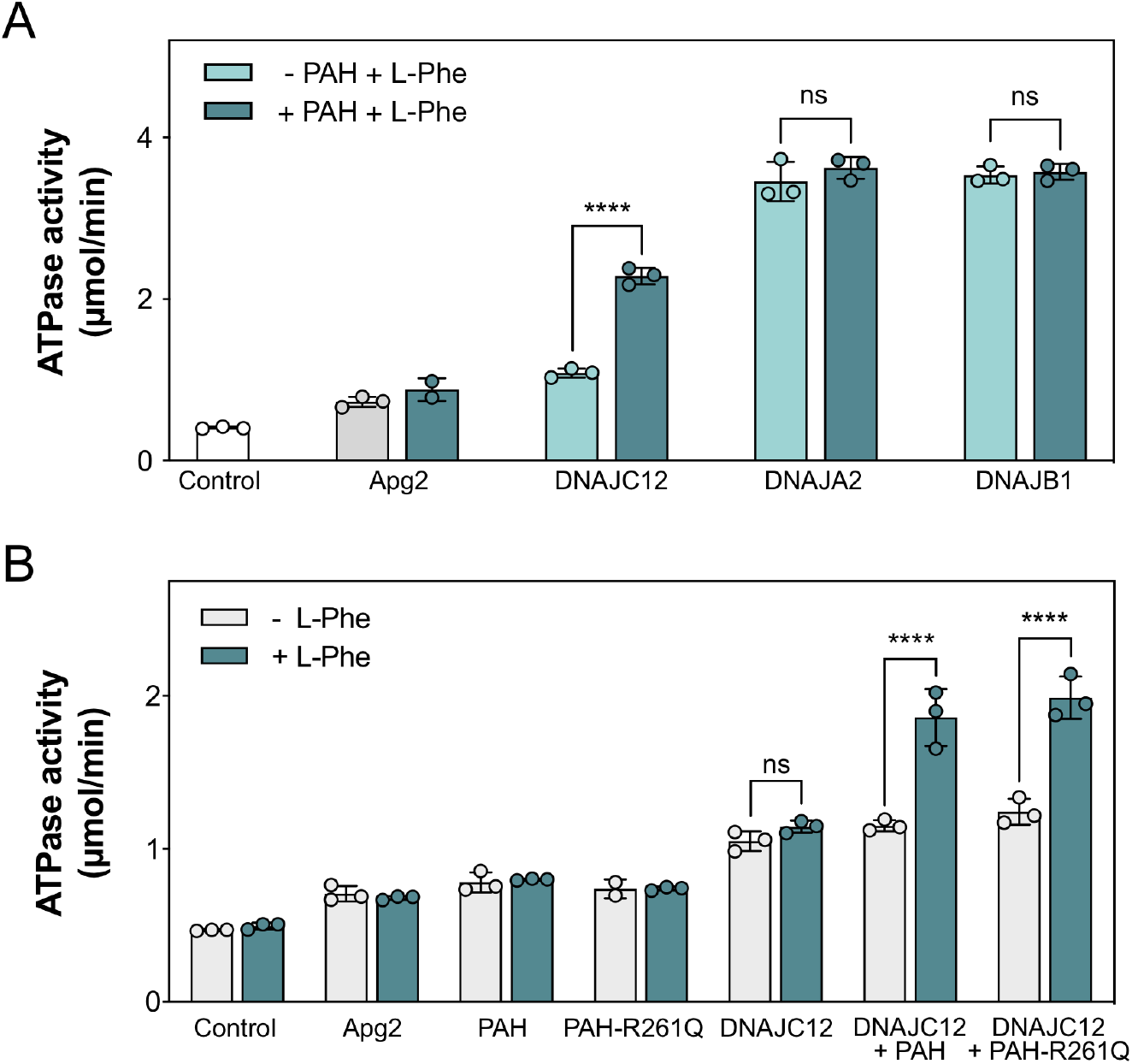
Hsc70 ATPase activity stimulation by DNAJC12 compared to model JDPs, in the presence and absence of L-Phe and/or PAH. **A. Stimulation of Hsc70 ATPase activity by JDPs with and without PAH, with 1 mM L-Phe**. Hsc70 ATPase activity was measured in the presence of DNAJC12 and the canonical JDPs DNAJA2 and DNAJB1, with (dark green) and without (light green) PAH. Data shown represent the mean ± SD for n=3 independent experiments. The ATPase activity recorded in the presence of PAH were compared to that recorded in the absence of the enzyme, by one-way ANOVA and Tukey’s post-hoc HSD test (****p<0.0001). **B. Stimulation of Hsc70 ATPase activity by DNAJC12 in the presence or absence of PAH, with and without 1 mM L-Phe**. The ability of DNAJC12 to stimulate Hsc70 ATPase activity was measured with and without PAH (WT or p.R261Q), with (dark green) or without (grey) of L-Phe. Data shown here represent the mean ± SD for n=3 independent experiments. The ATPase activity recorded with L-Phe were compared to their respective controls measured without L-Phe, by one-way ANOVA and Tukey’s post-hoc HSD test (****p<0.0001).

## DISCUSSION

An extensive network of molecular chaperones coordinate to maintain protein homeostasis. Thus, mutations in chaperones dysregulate these networks and may consequently affect the proteostasis of their client proteins, often leading to disease. Variants of the JDP *DNAJC12* cause HPA, dystonia, intellectual disabilities and neurotransmitter deficiencies, revealing the importance of this cochaperone in the regulation of the AAAHs^17-19^. The role of DNAJC12 in the maintenance of AAAH homeostasis has since been corroborated^20,22,23^, and characterization of the interaction between TH and DNAJC12 has recently improved the understanding of the determinants for DNAJC12-client binding^21^. However, little was known about the molecular mechanisms, as well as the regulatory and functional effects of PAH:DNAJC12 interaction. In this work, we identified the binding determinants for PAH:DNAJC12 complex formation as well as the consequences of this protein-protein interaction on the stability and activity of PAH.

Allosteric cooperativity is an important regulatory mechanism commonly found in metabolic and signaling pathways, and is usually accomplished through conformational changes induced by a ligand^29^. PAH is known to exist in different conformational states depending on the binding of its substrate, L-Phe^1,8,9^. At high substrate levels, L-Phe binding at the allosteric site in the ACT domains in the PAH-RDs leads to the consequent dimerization of these domains from opposite subunits in the tetramer^8,9^. The structure of the dimeric RDs with L-Phe bound at the interface has been solved using x-ray diffraction crystallography^10^, but the high resolution structure of the full-length substrate-bound PAH remains undetermined. In addition, the mechanism of substrate-induced activation remains unclear, but has been proposed to be initiated by L-Phe binding to the allosteric site in the RDs^9^. Substrate-induced dimerization of the PAH-RDs is proposed to increase the accessibility of the active site in the CD, as release of interactions between this domain and the autoregulatory region (residues 20-25) that covers the active site, initiates the transition of PAH from the precatalytic to the catalytic state^4,8^. Although this large activating conformational change increases PAH activity and stabilizes the tetramer^9,10^, the inherent dynamics necessary for maintaining high activity in this state may be challenging. Hence, increased interaction of DNAJC12 with activated PAH might have an implication on the overall stability of the enzyme in the activated state.

PAH:DNAJC12 complex formation was remarkably improved in the presence of L-Phe (Figs. 1A,D), indicating that the substrate-induced active conformation of PAH is likely a better physiological target for DNAJC12 than unactivated PAH. TH is not activated by its substrate L-Tyr or other amino acids, and therefore does not undergo the same conformational change as PAH, but already presents a comparable structure to activated PAH, with dimerized RDs^25,31^. Interestingly, DNAJC12 binds to unliganded TH effectively (K_D_=148 ± 18 nM)^21^, as it also does with activated PAH (K_D_=139 ± 19 nM). The full-length structures of the other two AAAHs, TPH1 and TPH2, are still undetermined and it is still uncertain whether L-Trp or other aromatic amino acids could activate these enzymes^32^, thus it is unknown whether DNAJC12 can readily bind to the unliganded forms of these enzymes as it does with TH.

When isolated from the rest of the protein, PAH-RDs exist as monomers that can be induced to form dimers with the addition of L-Phe^10^. By combining cryo-EM, crosslinking mass spectrometry, and site directed mutagenesis, we previously identified key residues in TH that are likely involved in complex formation with DNAJC12^21^. TH alanine variants at L141 and L145 exhibit improved affinity for DNAJC12 due to better packing, substantiating the involvement of these residues in hydrophobic interactions with the DNAJC12 CTD^21^. While the TH and PAH RDs share high structural homology (Fig. 6A), they differ in folding (Fig. 6B) and sequence (Fig. 6C). By aligning sequences and mapping the different secondary structures in the RDs of both proteins based on solved structures of human dimeric PAH RD (PDB 5FII)^10^ and TH RD (PDB 6ZVP)^25^, we found that the sequence that is involved in TH binding to DNAJC12 (^141^LAALL^145^)^21^ aligns with the rather similar sequence ^91^LTNII^95^ in PAH (Fig. 6C-D). The hydrophobic Leu residues involved in the interaction of TH with the DNAJC12 CTD^21^ appear preserved as Leu/Ile. This region is quite exposed in unactivated PAH (Fig. 6E; left), but it may be possible that the orientation of the RDs and steric hindrances imposed by the central CD+OD in this state prevents the efficient binding of DNAJC12, and explains the low-affinity interaction between DNAJC12 and unactivated PAH, as seen by the SEC, binding, DSF and aggregation assays without L-Phe. In the activated state, these residues are exposed in the same orientation observed in the analogous region in TH (Fig. 6D-E), and could explain why the affinity of DNAJC12 increases for activated PAH, to a similar level as previously reported for the TH:DNAJC12 interaction (K_D_= 148 nM)^21^. In addition, the 4:4 DNAJC12 monomer: PAH subunit stoichiometry suggests that two DNAJC12 monomers could bind to each RD dimer, potentially oriented in opposite directions to stabilize the interaction. This conformation may facilitate the retention of L-Phe within the RD, and potentially preserve some L-Phe to avoid depleting the amino acid during enzyme activation. However, the structural details and functional impact of PAH:DNAJC12 complex formation remain to be explored experimentally.

**Figure 6.**
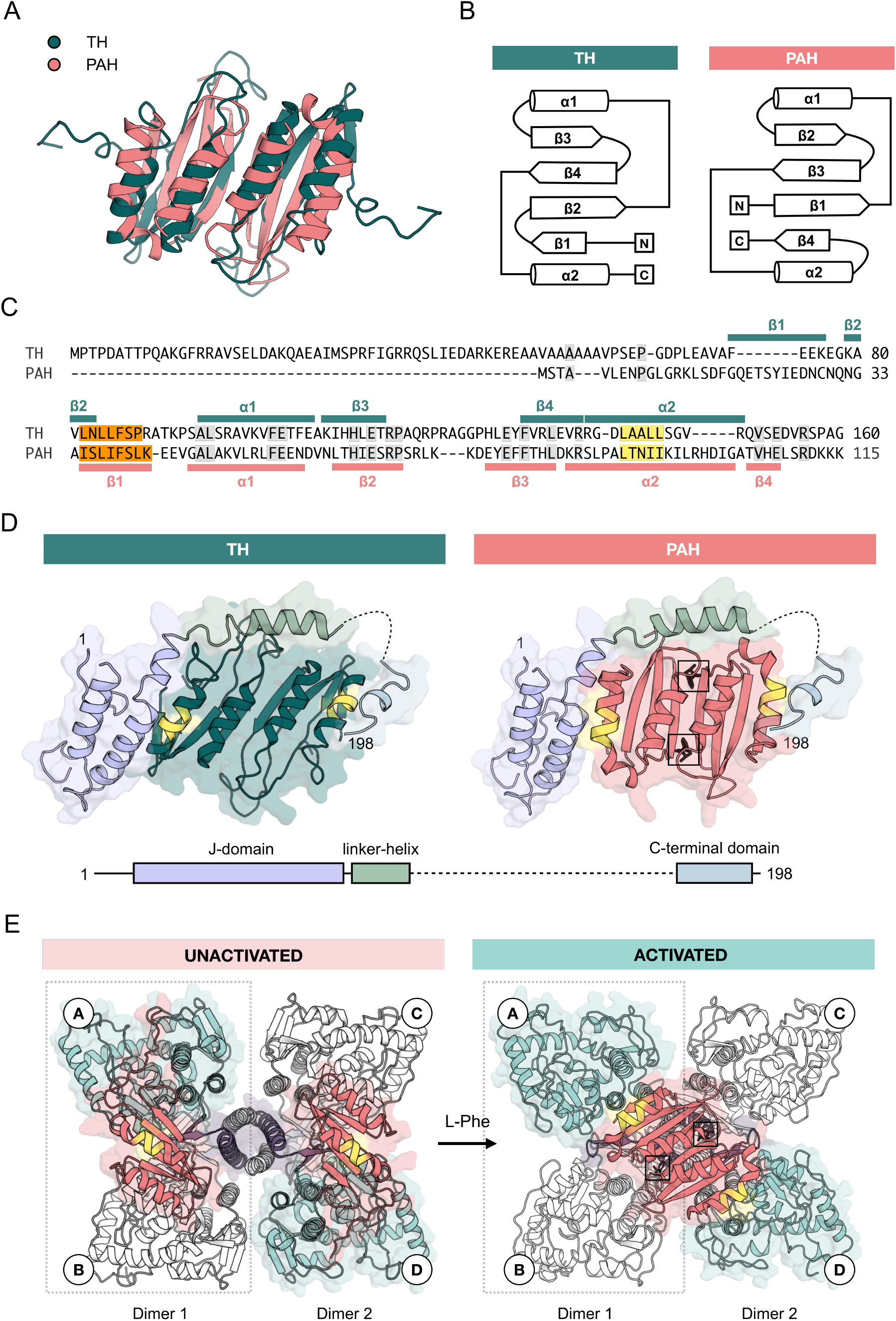
The modeled structure of activated PAH with the proposed PAH:DNAJC12 interaction region. **A. Structural comparison of the dimeric TH and PAH RDs**. Overlay of the human TH-RD solved by cryo-EM (PDB 6ZVP^25^;green) and of human PAH-RD with L-Phe solved by x-ray crystallography (PDB 5FII^10^; red) using PyMol. **B. Topology diagrams of the TH (green) and PAH RDs (red)**. The secondary structure elements of both the TH and PAH RDs are shown in the figure. **C. Sequence alignment of the TH and PAH RDs**. The sequence alignment, by ClustalOmega, and the secondary structures were mapped on the sequence based on the TH (PDB 6ZVP^25^;green) and PAH-RD (PDB 5FII^10^; red) structures. The region determined by Tai, et al.^21^ in the TH-RD (^141^LAALL^145^) that interacts with DNAJC12, and the equivalent sequence in PAH (^91^LTNII^95^) are highlighted in yellow, while the TANGO^39^-predicted residues in both TH (^82^LNLLFSP^88^)^21^ and PAH (^35^ISLIFSLK^42^), with high aggregation propensity, are highlighted in orange. **D. Model of PAH:DNAJC12 binding**. The dimeric PAH-RD crystal structure (PDB 5FII^10^;right) was aligned to the TH RD in the TH:DNAJC12 model that was developed based on cryo-EM, crosslinking-mass spectrometry and in silico analyses^21^ (left). The J-domain is highlighted in purple, the linker-helix in green and the client-binding CTD in blue. The proposed interaction region is colored in yellow in both models and the two L-Phe molecules bound in the dimeric PAH-RD are highlighted with a black box. **E. Model showing conformational changes in PAH induced by substrate binding**. Crystal structure of full-length unactivated human PAH (PDB 6HYC^4^; left) and a composite model of activated full-length PAH (right) made from the crystal structures of human CD+OD (PDB 2PAH) and human PAH RD (PDB 5FII^10^), with L-Phe bound on the dimeric interface (black box). The RD is colored in red, CD in blue and OD in purple. The site proposed to be involved in DNAJC12 interaction is in yellow.

In addition to their role as cochaperones of Hsp70, JDPs may directly protect regions in their client proteins that have a high propensity for aggregation by a holdase activity^33,34^ that has also been revealed for DNAJC12 with TH, preventing TH aggregation in vitro^21^. The aggregation prone residues ^82^LNLLFSP^88^ in the β1-strand in the TH-RD get covered upon DNAJC12 binding^21^. Similarly, the residues ^35^ISLIFSLK^42^ lying in the β1-strand of the PAH-RD are predicted by TANGO^39^ to have high propensity to engage in intermolecular cross-β interactions, and may also be covered by DNAJC12 binding. The binding of DNAJC12 to the dimeric RD could further contribute to prevent aggregation of PAH WT and PAH-p.R261Q by avoiding the dissociation of the dimers into monomers, a process that typically precedes aggregation^36,37^. L-Phe itself stabilizes PAH^10^ and facilitates RD dimer formation from different PAH dimers^8,35^, resulting in an additional tetramerization interface apart from the oligomerization domain (Fig. 6E). The strategic positioning of DNAJC12 could thus be especially valuable to stabilize aggregation-prone and unstable PAH variants such as PAH-p.R261Q. This substitution from Arg to Glu at position 261 is predicted to disrupt the interdimer interactions in PAH-p.R261Q and, consequently, the region surrounding the mutation may become prone to nonspecific intersubunit interactions^15^. Although DNAJC12 seems efficient in delaying the formation of PAH-p.R261Q aggregates in vitro, toxic amyloid-like PAH-p.R261Q aggregates have been identified in a mouse model^15^ despite the presence of WT DNAJC12. While our in vitro studies show that the binding stoichiometry between PAH and DNAJC12 is 1:1 per subunit, the mRNA expression of PAH mRNA has been reported to be approx. 20-fold higher than DNAJC12 in human liver samples^38^ (https://www.proteinatlas.org/). The disparity between expression levels could explain why DNAJC12 is unable to prevent PAH-p.R261Q aggregation in the mouse model. All in all, our in vitro experiments contribute to the understanding of DNAJC12 function and demonstrate the potential of increasing DNAJC12 expression or enhancing PAH:DNAJC12 interactions as a therapeutic strategy to stabilize and delay the aggregation of unstable PAH variants.

Previous in vitro characterizations have shown that the presence of L-Phe increases PAH activity^40^ as the enzyme transitions into its catalytic state^5,7^, showing half-maximal activity at approximately 190 μM (Fig. 3B), a concentration already considered within the range of mild HPA^3,41^. In this study, we found that DNAJC12 increases PAH activity at lower substrate concentrations and effectively reduces the *S*_0.5_ for L-Phe, without altering the positive cooperativity, to a level more relevant for healthy individuals, presenting an additional regulatory mechanism for PAH activity. The observed stabilization of PAH by DNAJC12 in the activated conformation could explain why the cochaperone increases PAH activity at lower L-Phe concentrations and why patients with DNAJC12 deficiency often present with HPA, as a consequence of this reduced activity in addition to the reduced stability^23^. Apart from these effects of DNAJC12 binding on PAH stability and activity, PAH:DNAJC12 complex formation seems to be essential for stimulation of Hsc70 ATPase activity by DNAJC12. Unlike better characterized JDPs such as DNAJA2 and DNAJB1, which are able to stimulate Hsc70 ATPase activity by themselves, without a client protein, DNAJC12 displays low stimulatory effect on its own, and synergistically stimulates Hsc70 ATPase activity when in complex with either TH^21^ or with PAH in an L-Phe dependent manner. These results suggest that the synergistic stimulation of Hsc70 ATPase activity by DNAJC12 may be preserved across its interactions with the four AAAH clients, notably in a dimerized RD conformation.

After several catalytic cycles, enzymes have a tendency to misfold or catalyze reactions inefficiently, often due to collateral damage from the reactions they facilitate^42^, and a catalysis-related loss of function due to generation of oxygen reactive species has indeed been shown for PAH^43,44^. The binding of DNAJC12 to activated PAH could thus enable Hsc70-mediated refolding or degradation to facilitate protein turnover. Overall, our findings suggest that DNAJC12 plays a crucial role in stabilizing PAH and enhancing its activity at lower L-Phe levels, and in maintaining PAH proteostasis, presenting the PAH:DNAJC12 interaction as a potential therapeutic target for HPA and PKU. Nevertheless, a long-term stabilization of PAH in the active state by DNAJC12 should be properly regulated, as it could result in the undesirable depletion of L-Phe, a crucial amino acid for protein synthesis.

## MATERIALS AND METHODS

### Plasmids

The pETMBP1a/*DNAJC12* plasmid encoding human WT DNAJC12, as well as the derivative plasmids encoding truncated DNAC12 variants DNAJC12(1-190) and DNAJC12(1-174 or Trp175Ter)^21^, as well as the pMAL-c5x/hPAH plasmid encoding WT PAH were previously available in the laboratory. Plasmids encoding PAH variants containing the RD (PAH(1-112)) or CD+OD (PAH(103-452)), and the disease-associated PAH variant PAH-p.R261Q were derived from the pMAL-c5x/hPAH plasmid (GenScript). The cDNAs of Apg2 (HSPH2), Hsc70 (HSPA8), DNAJA2 and DNAJB1 (Addgene) were cloned into the pE-SUMO vector (LifeSensors).

### Protein purification

Wild-type (WT) DNAJC12 and variants were expressed in *E. coli* BL21-CodonPlus(DE3)-RIL and purified as described by Tai, et al^21^. WT PAH and variants were purified from *E. coli* K12 TB1 cells (New England BioLabs) using a protocol previously described by Flydal, et al^4^. Recombinant proteins containing N-terminal His and SUMO tags (Apg2, Hsc70, DNAJA2, DNAJB1) were expressed in *E. coli* BL21 CodonPlus(DE3)-RIL or Rosetta™(DE3) as previously described by Cabrera et al.^45^ and Velasco-Carneros et al^46^.

### Analytical SEC

Samples containing 20 µM (subunit) PAH alone or with 20 µM DNAJC12 were prepared and analyzed on a Superdex™ 200 Increase (1.0 cm × 30 cm) column at a flow rate of 0.5 mL/min at 4°C, with a 20 mM HEPES, 200 mM NaCl pH 7 buffer or with the addition of 1 mM L-Phe.

### Purification of the PAH:DNAJC12 complex

The PAH:DNAJC12 complex was purified using SEC on a Superdex™ 200 Increase (1.0 cm × 30 cm) column by combining PAH and DNAJC12 proteins in a 4:8 PAH:DNAJC12 (subunits) molar ratio and collecting the fractions corresponding to the shifted peak (as observed in Fig. 1B). The run was done at a flow rate of 0.5 mL/min at 4 °C, with a 20 mM HEPES, 200 mM NaCl, 1 mM L-Phe, pH 7 buffer. Fractions corresponding to the PAH:DNAJC12 complex were pooled, collected, and concentrated and the protein concentration was determined by measuring the absorbance at 280 nm using the theoretical molar extinction coefficient A280 (1 mg.ml-1 cm-1) = 1.05.

### Native PAGE

Samples with 0.2 mg/mL DNAJC12, PAH or purified PAH:DNAJC12 complex in 20 mM Na-Hepes pH 7.0, 200 mM NaCl were diluted 1:1 with native PAGE Sample Buffer (Bio-Rad) and loaded into a 10% Mini-PROTEAN^®^ TGX Precast Protein Gel (Bio-Rad). The gel was run at 140 V for 3 hours at 4 °C with running buffer (25 mM Tris pH 8.3, 192 mM glycine), prior to visualization using 0.08 mM Coomassie brilliant blue G-250 (Bio-Rad) with 5 mM HCl. For samples where the samples contained either TH or PAH and different amounts of DNAJC12, 0.2 mg/mL PAH or TH with increasing amounts of DNAJC12 (4:1, 4:2 or 4:4).

### Immunoblotting

Proteins were transferred from the PAGE gel onto a polyvinylidene difluoride membrane (Bio-Rad) using the Transblot^®^ Turbo Transfer System (Bio-Rad) at 25 V for 3 min. The membrane was blocked with blocking solution (5% w/v skimmed milk powder, 1X Tris-buffered saline (TBS), 1% Tween 20). Primary antibody incubation was carried out overnight using a rabbit anti-DNAJC12 antibody (1:10000; ABCAM, cat. no. AB167425). The membrane was subsequently washed to reduce unspecific binding using washing buffer (1X TBS, 0.1% Tween 20) before incubation with goat anti-rabbit antibody (1:1000; Bio-Rad, cat. no. 1706515). The membrane was washed again prior to visualization using enhanced Luminita Immobilon^®^ Crescendo Western horse radish peroxidase substrate (Merck Millipore) and the ChemiDoc XRS+ System (Bio-Rad).

### Bio-layer interferometry

Bio-layer interferometry (BLI) was conducted using an OctetRED96 platform (FortéBio) with streptavidin (SA) biosensors (Sartorius). For PAH immobilization on the biosensor, PAH was biotinylated using EZ-Link™ NHS-PEG4-Biotin (Sigma Aldrich) according to the manufacturer’s instructions and diluted to 50 μg/mL in assay buffer (1x phosphate-buffered saline (PBS), 0.02% Tween-20, 0.5 mg/mL BSA) with or without 1 mM L-Phe. DNAJC12 was also prepared in the assay buffer at varying concentrations (0.01, 0.1, 0.5, 1, 2, and 4 μM) in the presence or absence of 1 mM L-Phe. The samples for were then dispensed into a 96-well flat-bottom polypropylene plate (Greiner), maintained at 25 °C, and agitated at 1000 rpm during the assay. Prior to each experiment, the biosensors were soaked in 1x PBS for 1-3 h, followed by equilibration in assay buffer for 30 s, PAH loading for 300 s, and re-equilibration in assay buffer for 120 s. The binding of DNAJC12 to immobilized PAH was monitored for 600 s, followed by a final equilibration in assay buffer for 300 seconds. Data were processed using the manufacturer’s software (Octet Data Analysis HT). Signals from the zero-concentration sample were subtracted from those obtained for each functionalized biosensor, plotted against the respective analyte concentrations, and fitted to a one-site binding (hyperbola) non-linear regression model to derive the KD values.

### SEC coupled with multi-angle light scattering

The sizes of PAH (2 mg/mL) and PAH:DNAJC12 complex (2 mg/mL) in 20 mM Na-Hepes pH 7.0, 200 mM NaCl were determined by SEC coupled with multi-angle light scattering (SEC-MALS). Samples were filtered using Corning^®^ Costar^®^ Spin-X^®^ (0.22 μm) plastic centrifuge tube filters (Merck) prior to sample application. The proteins were analyzed using a Superdex™ 200 Increase (1.0 cm × 30 cm) column connected to an iSeries LC-2050 (Shimadzu) HPLC system coupled to a RefractoMax 520 module (ERC GmbH) to measure the refractive index and determine concentration, and a mini-DAWN TREOS detector (Wyatt Technology) to measure light scattering. Data processing and molar mass estimation were performed using the Astra software (Wyatt).

### Differential scanning fluorimetry

Thermal denaturation curves of PAH alone or in the presence of DNAJC12, were obtained at different L-Phe concentrations using differential scanning fluorimetry (DSF). Samples with 1.92 µM (subunit) PAH alone or in presence of 3.84 µM DNAJC12 (4:8 PAH:DNAJC12 subunit ratio) were prepared with 2x FAS, 5x SYPRO Orange dye (Merck) in the presence or absence of L-Phe at varying concentrations (0-10 mM) prior to loading into a 384-well plate (Corning). The plate was heated step-wise from 25°C to 90°C at a heating rate of 2°C/min using a LightCycler 480 Real-Time PCR System (Roche Applied Science). The thermal denaturation of PAH was monitored by detecting the increase in SYPRO Orange fluorescence (λex = 465 nm, λem = 610 nm) upon its binding to exposed hydrophobic patches. Data analysis was done using HTSDSF Explorer^47^ to obtain the melting temperatures of the regulatory (*T*_m1_) and catalytic (*T*_m2_) domains.

### Dynamic light scattering

PAH aggregation was monitored in vitro by dynamic light scattering (DLS) using the Zetasizer Nano ZS instrument (Malvern Panalytical). Samples were prepared to contain 10 μM PAH (subunit) alone or with 10 μM DNAJC12 in 20 mM Na-Hepes pH 7.0, 200 mM NaCl, or withVan addition of 1mM L-Phe. The samples were centrifuged at 16,000 rpm for 5 min and filtered using Corning^®^ Costar^®^ Spin-X^®^ (0.22 μm) plastic centrifuge tube filters (Merck) prior to sample loading into UV cuvettes (Merck). Protein aggregation was analyzed by monitoring the change in the average particle size or Z-average value (diameters in nanometers; d.nm) of the protein samples over an 120-min period at 37 °C. Light scattering was measured every 2 min using a He-Ne laser at 683 nm with a fixed scattering angle of 173°. WT PAH samples containing L-Phe were incubated for an additional 30 min at 37 °C prior to collecting measurements.

### ATPase measurements

Steady-state ATPase assays were performed at 30 °C in 40 mM HEPES pH 7.6, 50 mM KCl, 5 mM Mg acetate, 2 mM DTT and an ATP regeneration-system (0.3 mM NADH, 3 mM phosphoenolpyruvate, 12.5 mg/mL pyruvate kinase, 0.017 mg/mL lactate dehydrogenase) in a final volume of 200 µL. Protein concentrations were 2 µM Hsc70, 0.4 µM Apg2, 1 µM JDPs, 1 mM PAH or PAH-p.R261Q. PAH and PAH-p.R261Q were initially incubated for 15 min at room temperature with 1 mM L-Phe, where indicated, followed by a further incubation for 15 min after addition of DNAJC12 or DNAJA2, or DNAJB1. The reaction was started after addition of Hsc70, Apg2 and 2 mM ATP (final concentration). ATP consumption was measured following the NADH decay by continuous measurement of the absorbance at 340 nm for 1 h using a Synergy HTX plate reader (BioTek) and transparent 96-well flat bottom plates (Sarstedt). The ATPase rates (µmol ATP/min) were calculated from the slopes of the A340 decay curves over the selected time intervals that showed a linear absorbance decline, using the extinction coefficient of NADH (ε340 = 6220 M^-1^ cm^-1^).

### PAH activity assay

PAH activity was assayed at 25 °C by continuous measurement of substrate production by assessing the increase in L-Tyr fluorescence intensity at an excitation wavelength of 274 nm and an emission wavelength of 304 nm using a Spark 20M microplate reader (Tecan). The standard reaction buffer containing 0.04 mg/mL catalase, 10 µM ammonium iron(II) sulphate, 1 mM L-Phe and 100 mM HEPES, pH 7.0 was added to sample wells on a black flat-bottom 96-well microplate (Greiner). PAH was added to a final concentration of 0.005 mg/mL with 0.05% BSA in the assay and preincubated for 3 minutes at 25 °C to be activated by L-Phe, without or with 0.01 mg/mL DNAJC12. The reaction was started by adding 75 µM BH_4_ with 2 mM DTT and the substrate production was measured in real-time for 15 min. To determine activity as a function of substrate, the concentration of L-Phe in the reaction mix was varied from 0-1 mM and BH_4_ was kept constant at 75 µM. Similarly, to determine activity as a function of the cofactor, the BH_4_ concentration was varied from 0-300 µM and L-Phe was kept constant at 1 mM. All given concentrations refer to the final concentration in a 50 µL reaction mixture. Fluorescence intensity was recorded for all enzyme activity measurements, and the reaction velocity was converted to enzyme activity (nmol Tyr/min/mg protein) using L-Tyr standards.

### Statistics and reproducibility

DSF dose response and PAH activity assay results were fitted to a nonlinear regression curve using the four-parameter dose-response model,

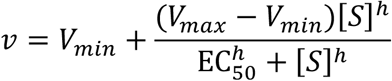

For the activity assays, *v* is the observed enzyme velocity, *V*_*min*_ and *V*_*max*_ are plateaus in the velocity, [*S*] is the substrate concentration and EC_50_ is the substrate concentration that gives half maximal velocity, and *h* is the Hill coefficient (GraphPad Prism 10.4.0). Values are given as the mean ± SD of three independent experiments done with technical triplicates. Accordingly, *v* is the observed enzyme *T*_m_, *V*_*min*_ and *V*_*max*_ are plateaus in the *T*_m_, while the EC_50_ represents the substrate concentration that gives half maximal stabilization, and are given as the mean ± SD of three independent experiments, for the DSF dose response assays. For the ATPase activity assays, at least three samples were prepared and measured prior to performing multiple comparisons using one-way analysis of variance (ANOVA) and a post-hoc Tukey HSD test. Results were considered to be significantly different when p<0.05.

## AUTHOR CONTRIBUTIONS

M.D.S.T., M.I.F. and C.F.D. purified PAH and DNAJC12, including all truncated forms and variants used in this work, and prepared the PAH:DNAJC12 complexes. M.D.S.T., G.G.-A., M.I.F. and C.F.D. carried out the biophysical analyses, while F.M. and T-A. K. performed biochemical experiments. M.D.S.T., M.I.F., J.P.K., F.M. and A.M. designed the experiments. All authors analyzed the data. A.M. managed the project and wrote the paper with main contributions from M.D.S.T. and corrections from all authors.

## ACKNOWLEDGEMENTS

We acknowledge the support of the Stiftelsen K.G. Jebsen (SKJ-MED-02; Center for Research in Parkinson’s Disease), The Neuro-SysMed Center (Research Council of Norway (RCN), Project No. 288164) and Fundació la Marató de TV3 (202012-31) to A.M., as well as the Meltzer Research Fund (Project 103578) to M.D.S.T. This research was also supported by grants PID2023-152081NB-I00 (MICIU/AEI/10.13039/501100011033 and FEDER, UE) and IT1745-22 (Basque Government) to F.M. We also acknowledge the Biophysics, Structural Biology and Screening (BiSS) core facility at the University of Bergen (UiB) and funding from NOR-OPENSCREEN (Grant No. 245922; Research Council of Norway (RCN). We are thankful for expert help from Dr. Lorea Velasco-Carneros, Kristine Kippersund Brokstad, Aamra Mahboob, Martine Therese Hopland and Sebastian Gonzalez Rodriguez and for access to the PNDdb (http://www.biopku.org).

